# Mechanical and chemical activation of GPR68 (OGR1) probed with a genetically-encoded fluorescent reporter

**DOI:** 10.1101/2020.04.02.022251

**Authors:** Alper D Ozkan, Tina Gettas, Audrey Sogata, Wynn Phaychanpheng, Jérôme J Lacroix

## Abstract

G-protein coupled receptor (GPCR) 68 (GPR68, or OGR1) couples extracellular acidifications and mechanical stimuli to G protein signaling and plays important roles in vascular physiology, neuroplasticity and cancer progression. Here, we designed a genetically-encoded fluorescent reporter of GPR68 activation called “iGlow”. iGlow responds to known GPR68 activators including fluid shear stress, extracellular pH and the synthetic agonist ogerin. Remarkably, iGlow activation occurred from both primary cilia-like structures and from intracellular vesicles, showing iGlow senses extracellular flow from within the cell. Flow-induced iGlow activation is not eliminated by pharmacological modulation of G protein signaling, disruption of actin filaments, or the presence of GsMTx4, a non-specific inhibitor of mechanosensitive ion channels. Genetic deletion of the conserved Helix 8, proposed to mediate GPCR mechanosensitivity, did not eliminate flow-induced iGlow activation, suggesting GPR68 uses a hitherto unkonwn, Helix8-independent mechanism to sense mechanical stimuli. iGlow will be useful to investigate the contribution of GPR68-mediated mechanotransduction in health and diseases.

## Introduction

G-protein coupled receptors (GPCRs) constitute the largest known family of membrane receptors, comprising at least 831 human homologs organized into 6 functional classes (A to F). They play essential roles in a wide range of biological functions spanning all major physiological systems including olfaction, energy homeostasis and blood pressure regulation. They also control embryonic development and tissue remodeling in adults. The biological significance of GPCRs is underscored by the fact that ~13% of all known human GPCRs represent the primary targets of ~34% of all pharmaceutical interventions approved by the Food and Drug Administration^1^.

GPCRs possess a conserved structure encompassing seven transmembrane helices and switch between resting and active conformations depending on the presence of specific physico-chemical stimuli. In addition to recognizing a vast repertoire of small molecules, such as odorants, hormones, cytokines, and neurotransmitters, GPCRs are also activated by other physico-chemical signals. These include visible photons^2^, ions^3^, membrane depolarizations^4–8^ and mechanical forces^9–12^. Once activated, GPCRs physically interact with heterotrimeric G-proteins (Gα, Gβ and Gγ), promoting the exchange of guanosine diphosphate (GDP) for guanosine triphosphate (GTP) on the Gα subunit. This process, called G-protein engagement, enables the GTP-bound Gα subunit and the Gβγ complex to dissociate from their receptor and activate downstream cellular effectors.

GPCRs often recognize more than one stimulus and interact with one or more Gα proteins amongst 18 known homologs, enabling them to finely tune downstream biological responses to a complex stimulus landscape^13^. One example of a GPCR exhibiting complex stimuli integration and pleiotropic G-protein signaling is GPR68, a class-A GPCR first identified in an ovarian cancer cell line and hence initially named Ovarian cancer G-protein coupled Receptor 1, or OGR1^14^. GPR68 is expressed in various tissues and is often up-regulated in many types of cancers^9,15^. Although sphyngophosphorylcholine lipids were proposed to act as endogenous GPR68 ligands^16,17^, one of these two studies has been subsequently retracted^18^. It is now well-established that GPR68 is physiologically activated by extracellular protons^19^, a property shared with only three other GPCRs to date (GPR4, GPR65 and GPR132). As reported for several GPCRs, GPR68 is also activated by endogenous mechanical stimuli, such as fluid shear stress and membrane stretch^9,10,20^. This mechanosensitivity enables GPR68 to mediate flow-induced dilation in small-diameter arteries^9^. GPR68 can engage Gα_q/11_, which increases cytosolic concentration of calcium ions ([Ca^2+^]_cyt_) through phospholipase C-β (PLC-β), as well as Gα_s_, which increases the production of cyclic adenosine monophosphate (cAMP) through adenylate cyclase activation. Interestingly, the synthetic GPR68 agonist ogerin increases pH-dependent cAMP production by GPR68 but reduces pH-dependent calcium signals, suggesting that ogerin acts as a biased positive allosteric modulator of GPR68^21^. Interestingly, ogerin suppresses recall in fear conditioning in wild-type but not GPR68^-/-^ mice, suggesting a role of GPR68 in learning and memory^21^. Hence, although the contribution of GPR68 to vascular physiology has been relatively well-established, its roles in other organs such as the nervous and immune systems remain unclear. It is also unclear whether the molecular mechanism underlying GPR68 mechanosensitivity is shared with other mechanosensitive GPCRs.

A genetically-encoded fluroescent reporter of GRP68 activation would facilitate these investigations. GPCR activation is often monitored using Fluorescence Resonance Energy Transfer (FRET) or Bioluminescence Resonance Energy Transfer (BRET)^11,20,22^. However, FRET necessitates complex measurements to separate donor and acceptor emissions whereas BRET often requires long integration times and sensitive detectors to capture faint signals. In contrast, recent ligand-binding reporters engineered by fusing GPCR with a cyclic permuted Green Fluorescent Protein (cpGFP) have enabled robust and rapid *in vitro* and *in vivo* detection of GPCR activation by dopamine^23,24^, acetylcholine^25^ and norepinephrine^26^ using simple intensity-based fluorescence measurements. Here, we borrowed a similar cpGFP-based engineering approach to create a genetically-encodable reporter of GPR68-mediated signaling. We call it indicator of GPR68 activation by f<u>l</u>ow and l<u>ow</u> pH, or iGlow.

## Results

### iGlow senses flow-induced GPR68 activation

We designed iGlow by borrowing a protein engineering design from dLight1.2, a genetically-encoded dopamine sensor^23^. In dLight1.2, fluorescence signals originate from a cyclic permuted green fluorescent protein (cpGFP) inserted into the third intracellular loop of the DRD1 dopamine receptor. We generated iGlow by inserting cpGFP into the homologous position in the third intracellular loop of human GPR68, flanking cpGFP with the same N-terminal (LSSLI) and C-terminal (NHDQL) linkers as in dLight1.2 (**Figure 1A, Supplementary Figure 1 and Supplementary Table 1**). To study iGlow under flow, we transiently expressed iGlow into HEK293T cells and seeded them onto microscope-compatible flow chambers, allowing us to apply desired amount of fluid shear stress (FSS) by circulating Hank’s Balanced Salt Solution (HBSS, pH 7.3) using a computer-controlled peristaltic pump while recording fluorescence with an inverted epifluorescence microscope.

**Figure 1:**
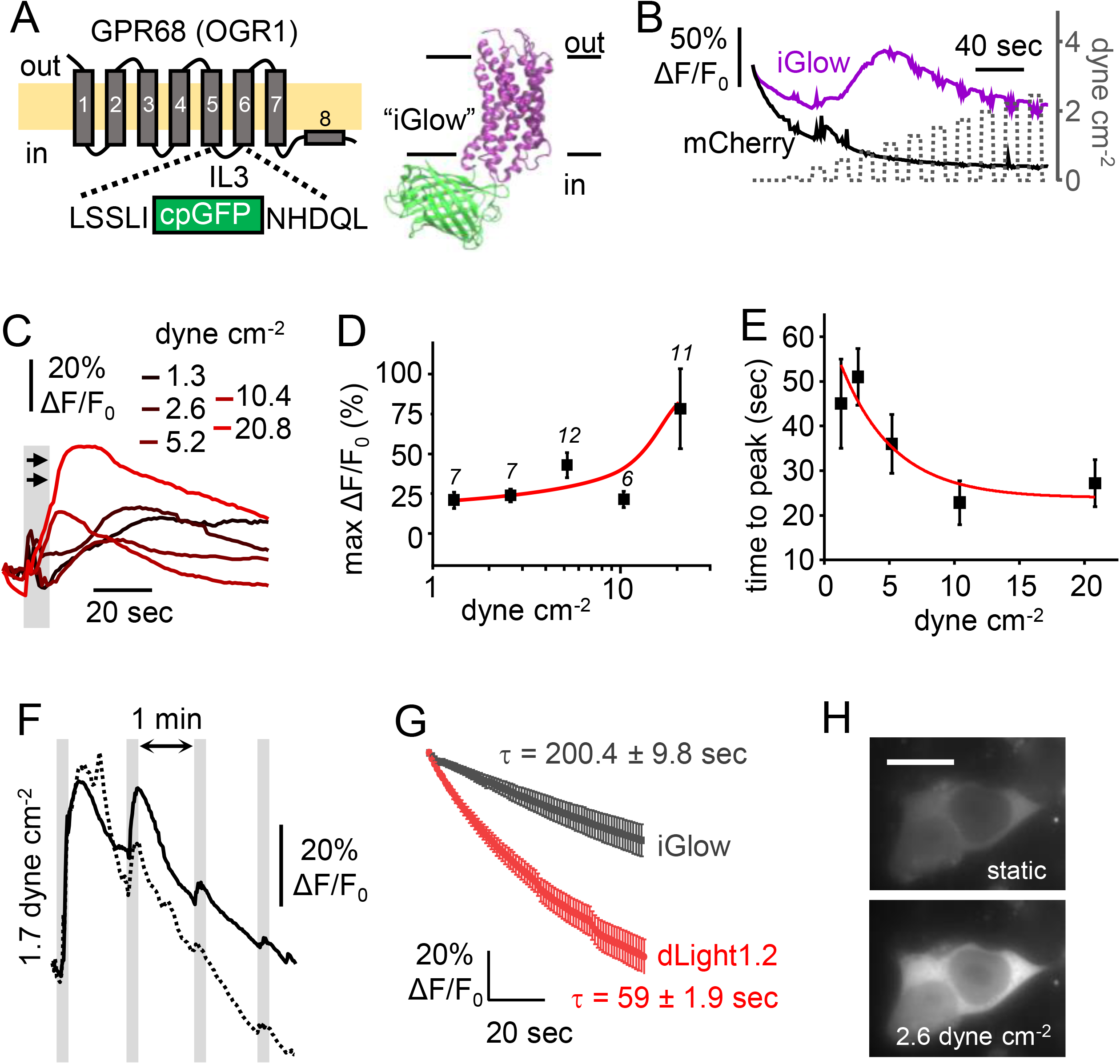
Design and characterization of iGlow. (**A**) *Left*: iGlow was designed by inserting a cpGFP cassette (green) into the third intracellular loop (IL3) of GPR68 (purple). *Right*: structural model of iGlow generated using the Molecular Operating Environment (MOE) software from the crystal structure of cpGFP (PDBID: 3O77, green) and a structural model of GPR68 (purple) generated by Huang et al.^21^. (**B**) Fluorescence time-courses from a cell co-expressing iGlow (purple trace) and mCherry (black trace) in response to intermittent shear stress pulses (10 sec on, 10 sec off) of incrementally-increased amplitudes. (**C**) iGlow fluorescence signals evoked by single shear stress pulses (grey bar) of indicated amplitude. (**D**) Scatter plot showing the maximal ΔF/F_0_ values produced by iGlow using data from (C). Numbers above data indicate n values. Red line = trend line (**E**) Time to peak values plotted as function of the shear stress amplitude. Red line = mono-exponential fit (R^2^ = 0.81). (**F**) Fluorescence time course from two independent iGlow expressing cells (solid and dashed lines) obtained with repeated shear stress pulse (1.7 dyne cm^-2^) with 1 min recovery. (**G**) Averaged fluorescence photobleaching obtained from cells expressing iGlow (grey, n = 10) or dLight1.2 (red, n = 9) exposed to the same illumination protocol. τ = fitted exponential time constants. (H) Epifluorescence images showing iGlow fluorescence in static or flow condition (bar = 10 μm). Error bars = s.e.m.

In order to accurately determine the amplitude of shear stress applied inside the flow chambers, we first compared shear stress values determined by multiplying the average flow-rate by a coefficient provided by the manufacturer to values calculated by particle velocimetry measurement (**see Methods and Supplementary Figure 2**). We observed that increasing flow rate tends to increase fluctuations of velocity measurements, yielding a poor linear fit between flow rate and shear stress (R^2^ = 0.19). We attribute these fluctuations to the peristaltic nature of the flow, which inevitably produces fluctuations in instantaneous flow rate. Nevertheless, the fitted slope value obtained from velocity measurements was near the manufacturer’s calibration (1.75 ± 0.65 dyne mL cm^-2^ min^-1^ vs. 1.316 dyne mL cm^-2^ min^-1^). Hence, we used the manufacturer’s calibration to determine the amplitude of shear stress in all our experiments.

To rapidly explore the flow sensitivity of iGlow, we stimulated cells co-expressing iGlow and mCherry with an “escalating” stimulation protocol (10 sec on, 10 sec off, using incrementally increased FSS amplitudes). This protocol produced robust and transient increases in green, but not red, fluorescence intensity, indicating green fluorescence changes did not result from imaging artifacts (**Figure 1B**). To rule out direct modulation of cpGFP fluorescence by FSS independently of GPR68, we first tested whether dLight1.2 responds to FSS. Our data show that dLight1.2 produces small but nevertheless detectable fluorescence changes upon FSS, perhaps due to the endogenous sensitivity of dopamine receptor homologs to shear stress^27^ (**Supplementary Figure 3A-B**). We next measured green fluorescence from two other membrane proteins containing cpGFP which are not known (or anticipated) to exhibit mechanosensitivity: the voltage indicator ASAP1^28^ and Lck-cpGFP, a cpGFP fused to the membrane-bound N-terminal domain of the Lymphocyte-specific kinase (Lck)^29^. In these cpGFP-containing proteins, no fluorescence change other than photobleaching-induced decay occurred, even upon high flow conditions (> 10 dyne cm^-2^), confirming FSS-induced signals from cpGFP inserted into iGlow require the presence of GPR68 (**Supplementary Figure 3C-E**).

We next determined the dynamic range of iGlow fluorescence by exposing cells to single FSS pulses ranging between 1.3 and 20.8 dynes cm^-2^ (**Figure 1C**). We used the amplitude of the fluorescence peaks, calculated as max ΔF/F_0_, to quantify the amount of iGlow activation. The peak amplitude gradually increases over the tested shear stress range, the largest increase being a nearly 3-fold increase between 10.4 and 20.8 dynes cm^-2^ (approximately 25% to 75%), which matches well the range of GPR68 activation measured with calcium imaging^9^ (**Figure 1D**). In addition, the delay between mechanical stimulation and peak fluorescence (time to peak) was negatively correlated with the amplitude of the FSS pulses (Pearson’s correlation coefficient = −0.77), stronger pulses producing shorter time to peak values. The relationship between pulse amplitude and time to peak is not linear and was best fitted by an exponential function (R^2^ = 0.81, red trace in **Figure 1E**).

Next, we performed a multiple-pulse protocol with 1 min interval between each pulse to measure the repeatability of iGlow signals. Our data indicate that, although subsequent pulses induce subsequent fluorescence peaks, the amplitude of these secondary peaks was lower than the first one (**Figure 1F**). This signal loss may be due to excessive photobleaching, receptor desensitization, or to an adaptation of the cell to repeated shear stress. Although we applied a photobleaching correction for all recordings (except for the escalating stimulation, Fig 1B), we sought to directly evaluate iGlow photobleaching by fitting spontaneous iGlow and dLight1.2 fluorescence decay in absence of stimulation to an exponential function. Our data indicate faster photobleaching in dLight1.2 compared to iGlow (time constants of 59.0 ± 1.9 sec, R^2^ = 0.997; and 204.4 ± 9.8 sec, R^2^ = 0.999; respectively) (**Figure 1G**).

We noticed that, in many cells, both baseline iGlow fluorescence (i.e. before stimulation) and FSS-induced signals originate from the whole cell (**Figure 1H**), in contrast to other tested cpGFP-containing membrane proteins showing exclusive membrane fluorescence (**Supplementary Figure 3A and 3D**). This prompted us to investigate the cellular localization of iGlow. To this aim, HEK293T cells were transfected with plasmids encoding iGlow and a red fluorescent marker of actin (LifeAct-mScarlet), or one of two red fluorescent proteins fused with a C-terminal CAAX motif (dsRed-CAAX and HcRed-CAAX) essential for membrane trafficking of RAS proteins^30^. Live-cell confocal fluorescence imaging shows that iGlow is sparsely present in intracellular compartments in some cells, while in other cells, iGlow preferentially co-localizes with cortical actin filaments and is often found in peripheral cellular structures resembling primary cilia (**Figure 2A**). Furthermore, epifluroescence imaging of cells in static or flow conditions shows that iGlow activation occurs from both the plasma membrane and/or cilia-like structures (**Figure 2B**) as well as from subcellular vesicular compartments (**Figure 2C, red arrows**).

**Figure 2:**
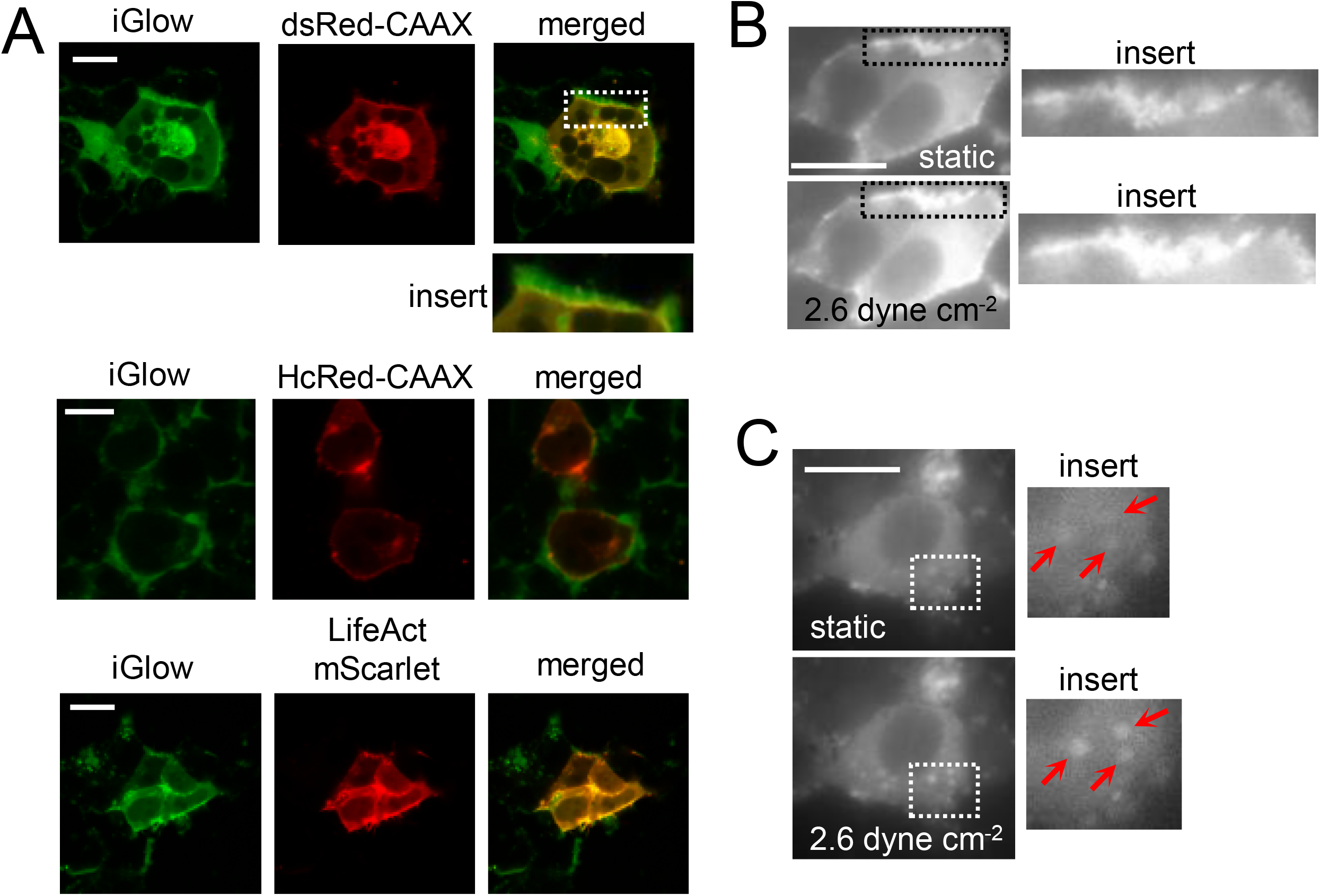
iGlow activation occurs at both plasma and subcellular membranes. (**A**) Live-cell confocal images of cells co-expressing iGlow and dsRed-CAAX (top), HcRed-CAAX (middle), or LifeAct-mScarlet (bottom). (**B**-**C**) Epifluorescence images of iGlow-expressing cells before (static) or during a shear stress pulse of 2.6 dyne cm^-2^. Insert locations are indicated by dashed boxes. In all panels, horizontal bars = 10 μm.

### iGlow senses chemical GPR68 activation

Next, we investigated the ability of iGlow to sense ogerin and extracellular acidifications. For this, we first acutely perfused iGlow-expressing cells with a physiological solution containing 10 μM ogerin or a vehicle control and measured iGlow fluorescence (**Figure 3A**). As positive control, we measured dLight1.2 response to 10 μM dopamine (**Figure 3B**). We observed rapid and transient iGlow activation events that were significantly higher than the vehicle control (Mann-Whitney U-test p-value = 0.0271) (**Figure 3C**). We also confirmed dLight1.2 sensitivity to dopamine, albeit with lower responses than reported by Patriarchi et al. (approximately 50% vs. >300%)^23^. We attribute this discrepancy to the fact that we collected dLight1.2 signals from whole cell pixels rather than membrane pixels and with epifluorescence instead of confocal fluorescence (which increases signal-to-noise ratio). The time to peak values of ogerin-induced iGlow signals were also significantly shorter when stimulating iGlow with 10 μM ogerin as compared to a 20.8 dyne cm^-2^ FSS pulse (Mann-Whitney U-test p-value = 0.0307) (**Figure 3D**).

**Figure 3:**
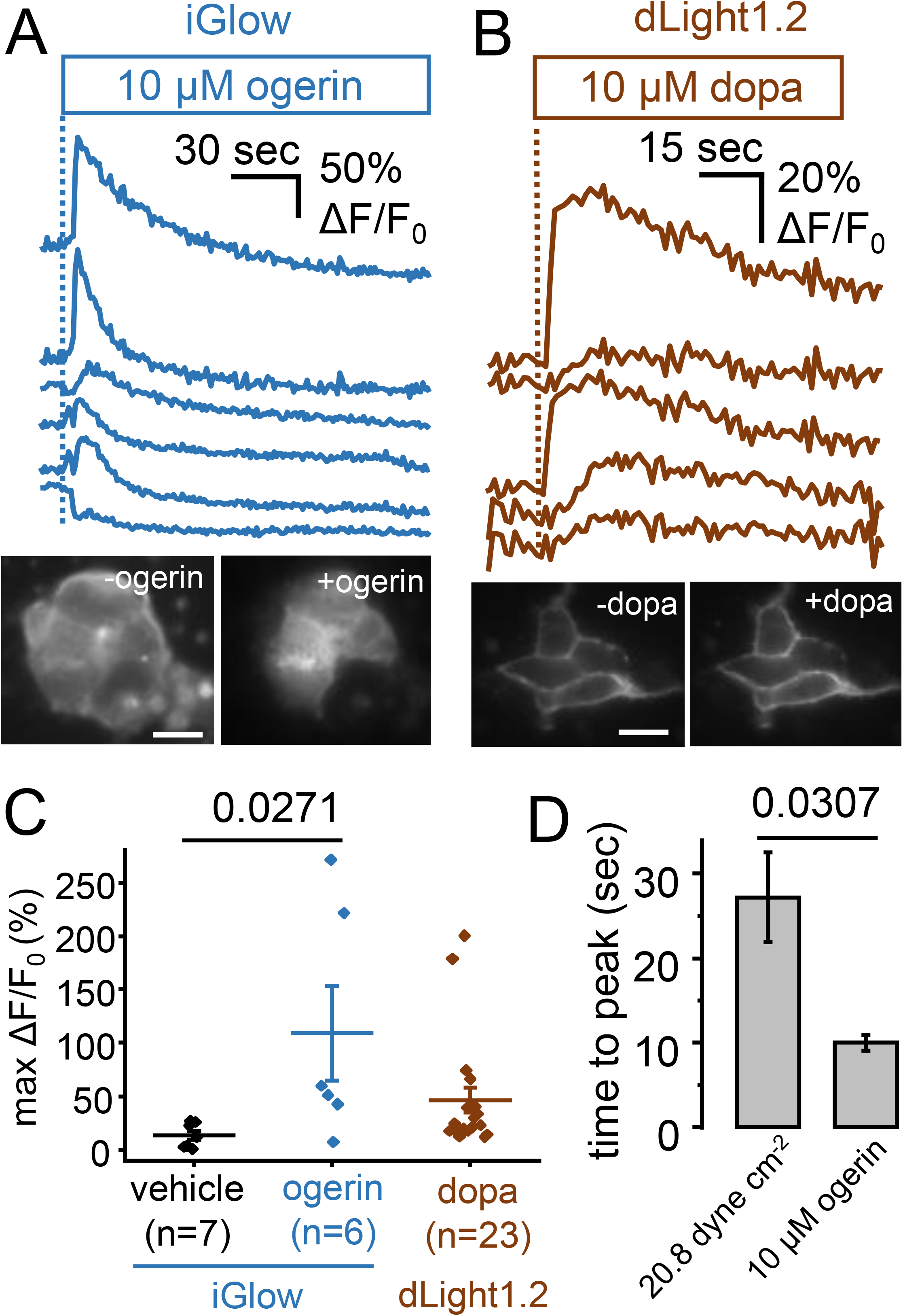
Chemical activation of iGlow with ogerin. (**A**) *Top*: Fluorescence time-course of iGlow expressing cells in response to acute perfusion with 10 μM ogerin. *Bottom*: images of cells before and after ogerin perfusion. (**B**) *Top*: Fluorescence time-course of dLight1.2 expressing cells in response to acute perfusion with 10 μM dopamine (dopa). *Bottom*: images of cells before and after dopamine perfusion. (**C**) Scatter plots showing max ΔF/F_0_ values obtained from iGlow expressing cells perfused with ogerin (blue dots, n = 6) or a vehicle control (black dots, n = 7) or from dLight1.2 expressing cells perfused with dopamine (brown dots, n = 23). (**D**) Comparison of time to peak values obtained from iGlow expressing cells exposed to a high amplitude shear stress pulse or 10 μM ogerin. Numbers above plots in (C) and (D) indicate exact p-values from Mann Whitney U-tests (C). In panels (A-B), horizontal bars = 10 μm. Error bars = s.e.m.

We next manually perfused iGlow-expressing cells with an acidic physiological solution, decreasing extracellular pH from 7.3 to 6.5, or with a vehicle control maintaining a neutral external pH. iGlow produced robust and transient signals that were larger at pH 6.5 (**Figure 4A-B**). In many cases, iGlow also responded modestly to the vehicle control, suggesting that iGlow is activated by the shear stress producing by the perfusion. We also noticed this perfusion artefact in the ogerin experiments (**Figure 3C**). Hence, to avoid fluctuations due to the unpredictable shear stress component of manual perfusion, we stimulated iGlow-expressing cells with a single FSS pulse of 2.6 dyne cm^-2^ using a physiological solutions at pH of 8.2 (n = 28), 7.3 (n = 24) or 6.5 (n = 10), starting from pH 8.2 to reduce baseline activation of GPR68^9^ (**Figure 4C**). Our data show that, on average, iGlow signals tend to be inversely correlated with pH, with peak fluorescence values significantly different among all treated groups (one-way ANOVA p-value 0.0090) (**Figure 4D-E**). In line with the effect of extracellular acidification on signal amplitude, the mean time to peak value was also significantly shorter at pH 6.5 compared to 7.3 (Student’s T-test p-value = 0.0076), while no statistically significant difference was observed between pH 7.3 and 8.2 (Student’s T-test p-value = 0.2570) (**Figure 4F**).

**Figure 4:**
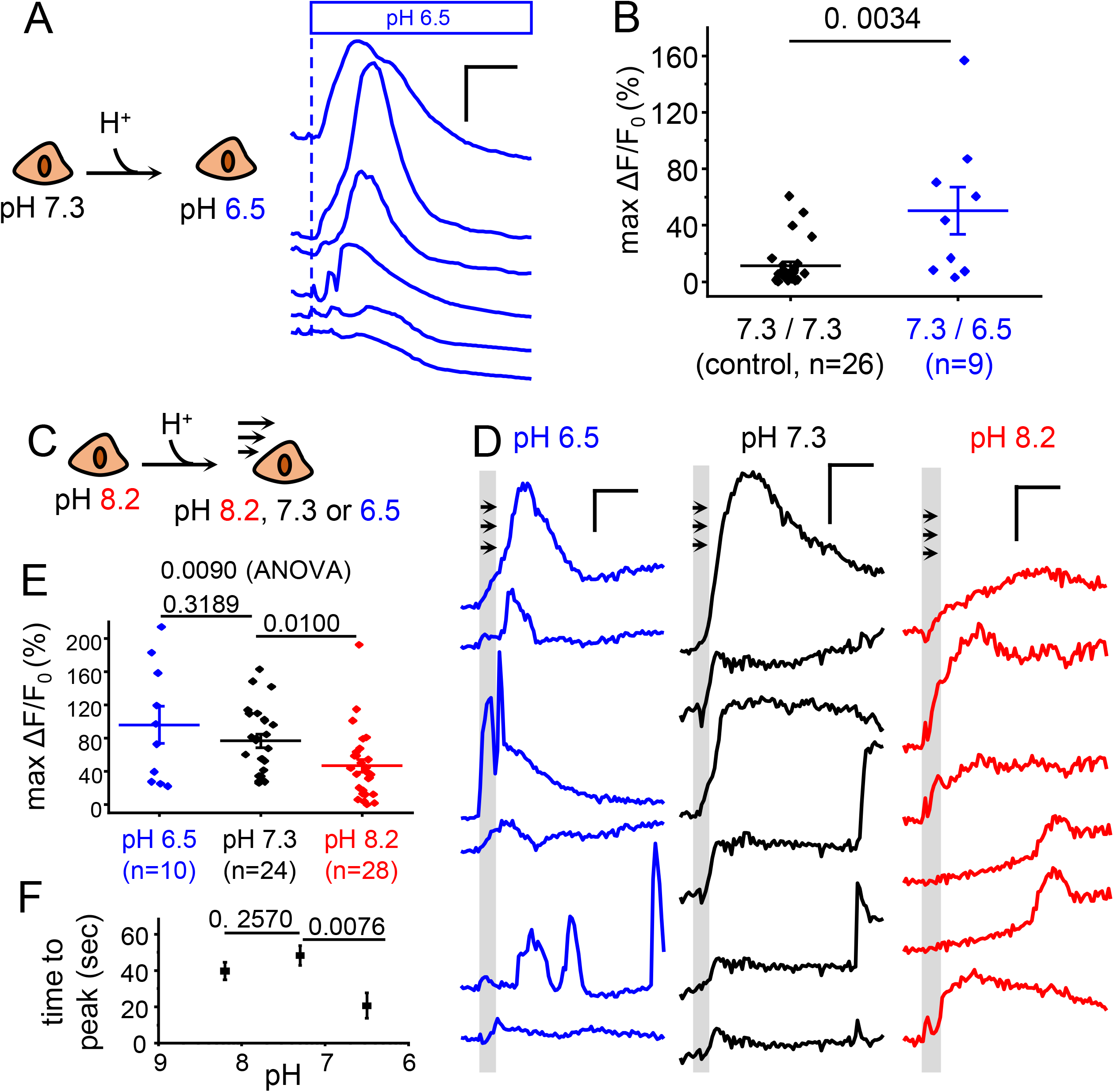
Modulation of iGlow by extracellular pH. (**A**) Examples of fluorescence time course from individual iGlow-expressing cells exposed to acute extracellular acidification from pH 7.3 to pH 6.5. (**B**) Scatter plots showing max ΔF/F_0_ values from data obtained from (A) (7.3 / 6.5, n = 9) and from control experiments with no pH change (7.3 / 7.3, n = 26). (**C**) iGlow expressing cells were incubated at pH 8.2 and exposed to a single shear stress pulse using extracellular solution of indicated pHs. (**D**) Representative fluorescence time course of single iGlow-expressing cells from experiment depicted in (C). Vertical bars = 50% ΔF/F_0_, horizontal bars = 20 sec. (**E**) Scatter plots showing max ΔF/F_0_ values from experiments depicted in (C). (**F**) Plot showing the mean time to peak values as function of extracellular pH from data shown in (E). Numbers above plots indicate exact p-values from Mann Whitney U-test (B), Student’s T-tests (E-F), or one-way ANOVA (E). Error bars = s.e.m.

### Flow-induced iGlow activation is independent of G proteins

To test if iGlow signals depend on specific interactions between GPR68 and its regulatory cytoplasmic proteins, we stimulated iGlow with a 2.6 dyne cm^-2^ FSS pulse in cells pre-treated one of several pharmacological agents. We used GTP-γ-S, a non-hydrolyzable GTP analog which prevents Gα protein association to all GPCRs; NF449, a GDP→GTP exchange inhibitor which selectively prevents Gαs dissociation from its receptor^31^; and BIM-46187, a non-specific GDP→GTP exchange Gα inhibitor^32^. We also tested CMPD101, an inhibitor of G-protein Receptor kinases 2/3 (GRK2/3) (**Figure 5A**). Most treatments did not significantly change the mean peak fluorescence nor the time to peak values (**Figure 5B-D**). One exception was the GTP-γ-S treatment, which showed a significant increase in mean max ΔF/F_0_ from 66 ± 6 % (control) to 100 ± 9 % (Student’s T-test p-value = 0.0028). We also observed that dopamine-induced dLight1.2 signals were not significantly affected by any of these pharmacological treatments (one-way ANOVA p-value = 0.5441) (**Supplementary Figure 4**), as expected since dLight1.2 was shown to apparently block the ability of the dopamine receptor to interact with G-proteins^23^.

**Figure 5:**
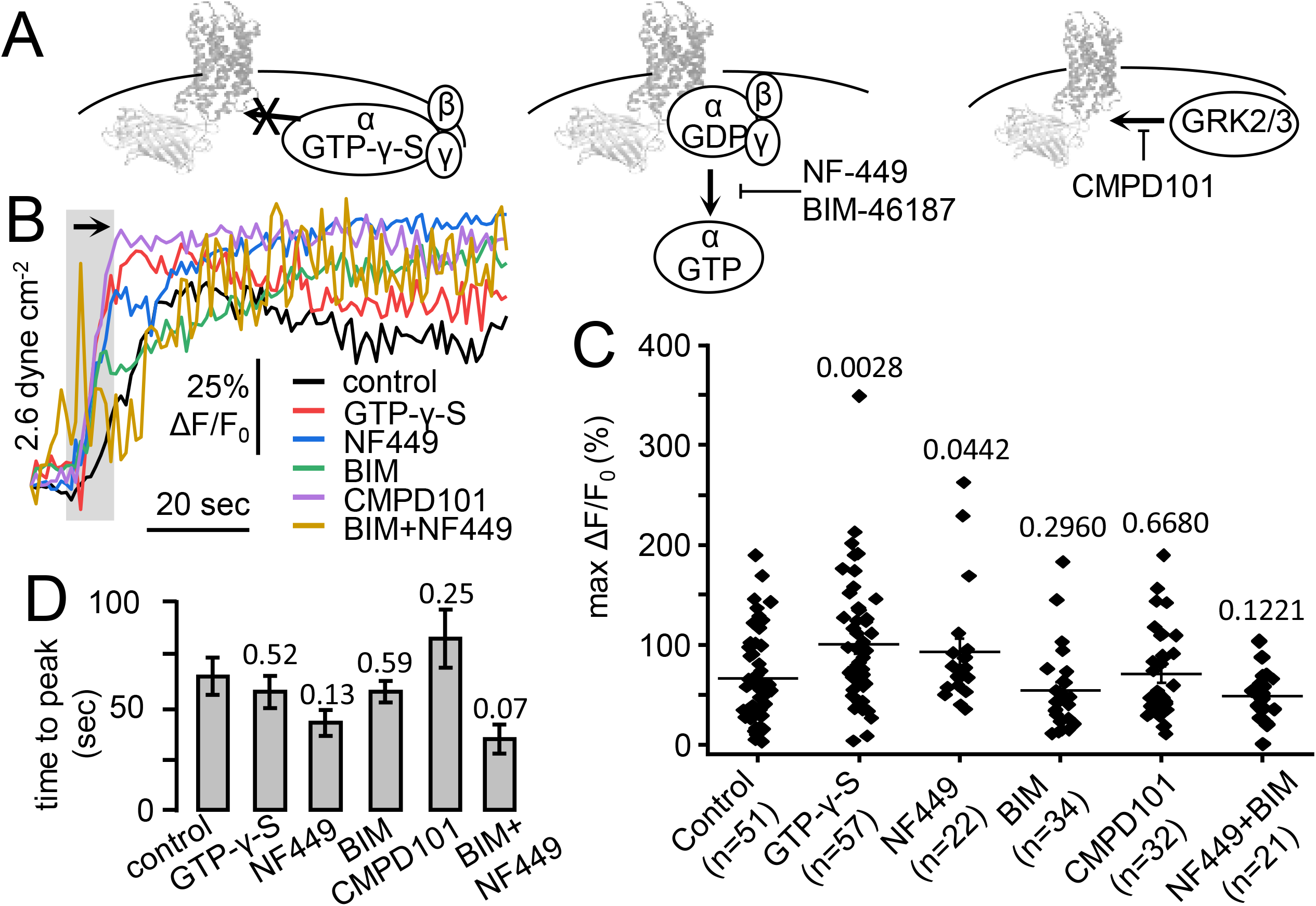
Flow-induced iGlow signals are not abolished by pharmacological modulation of downstream G-protein signaling. (**A**) Expected effects of pharmacological treatments on protein-protein interactions between iGlow, Gα proteins and GRK2/3 kinases. (**B**) Representative fluorescence time-course of iGlow-expressing cells treated with 0.2 mM GTP-γ-S (red trace), 20 μM NF-449 (blue trace), 20 μM BIM-46187 (BIM, green trace), 10 μM CMPD101 (purple trace) or a vehicle control (black trace) and exposed to an acute shear stress pulse. (**C**) Scatter plots showing the max ΔF/F_0_ values obtained following shear stress stimulation in cells treated with GTP-γ-S (n = 33), NF-449 (n = 20), BIM-46187 (BIM, n = 21), CMPD101 (n = 17) or a vehicle control (n =25). (**D**) Histograms showing the mean time to peak values from data obtained in (C). Numbers above plots or bars in panels (C-D) indicate exact p-values from Student’s T-tests between control and treated samples. Error bars = s.e.m.

### Flow-induced iGlow activation is robust

We next sought to determine whether iGlow signals depend on the integrity of the actin cytoskeleton. To this aim, we transfected HEK293T cells with LifeAct-mScarlet to monitor real-time actin disorganization upon treatment with 20 μM cytochalasin D (CD20), an inhibitor of actin polymerization. Actin filaments were completely disorganized after about 20 min (**Figure 6A**). Since we observed visible cell death after a one-hour 20 μM CD treatment, we monitored iGlow’s response to FSS immediately after 20 min incubation. At the shear amplitude of 2.6 dyne cm^-2^, iGlow produced fluorescence signals similar to untreated cells, even upon increasing CD concentration to 50 μM CD (one-way ANOVA p-value = 0.6825) (**Figure 6B-C**). These results show that, at this shear stress amplitude, the actin cytoskeleton is not necessary for shear stress sensing by iGlow.

**Figure 6:**
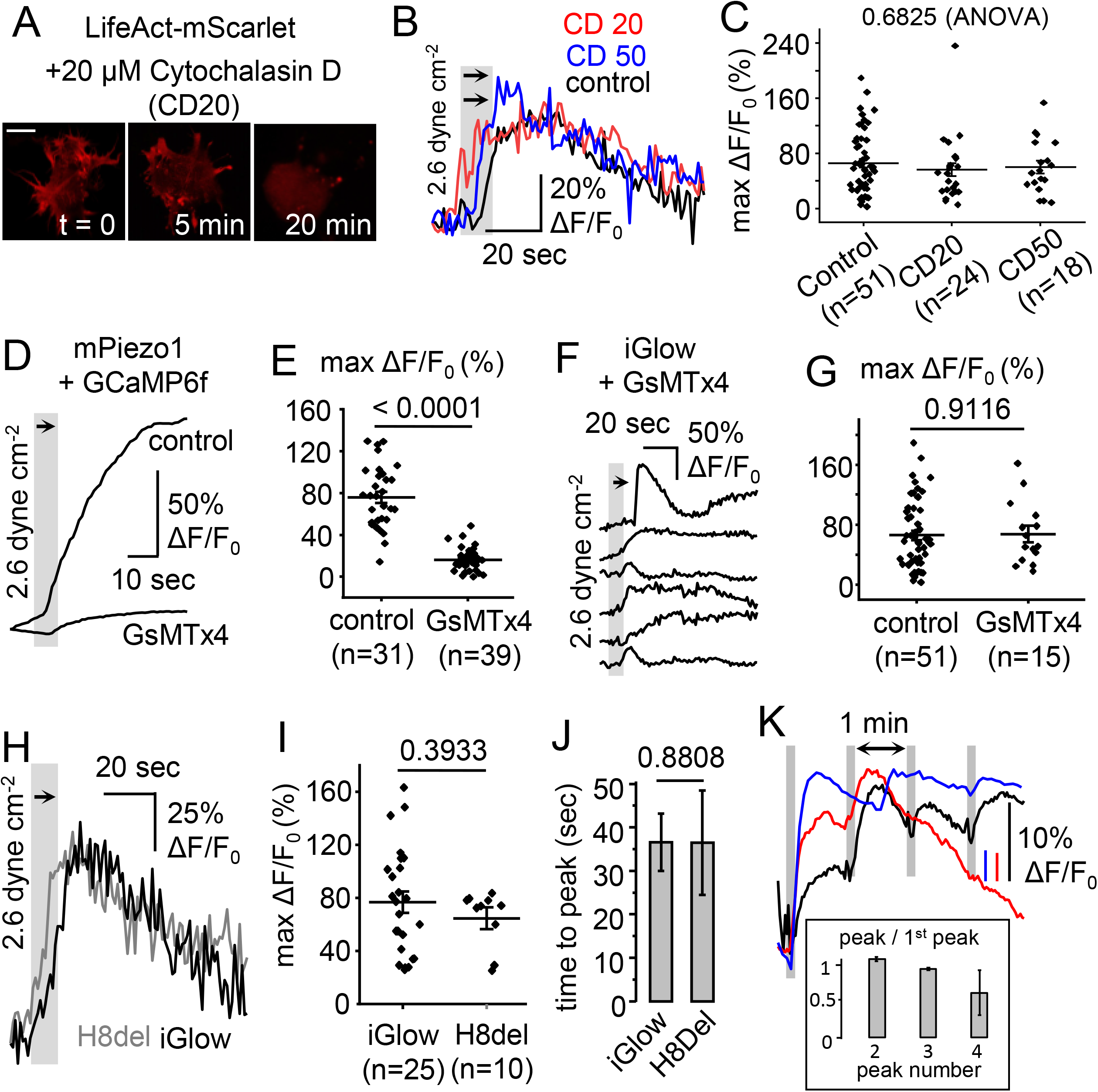
Flow-induced iGlow signals are not abolished by experimental manipulations known to modulate mechanosensitivity of ion channels or GPCRS. (**A**) Confocal images of a LifeAct-mScarlet-expressing cell at indicated times following incubation with 20 μM Cytochalasin D (CD) (bar = 10 μm). (**B**) Representative fluorescence time-course of iGlow from cells incubated for 20 min with 20 μM CD (red), 50 μM CD (blue) or a control solution (black) and exposed to a shear stress pulse. (**C**) Scatter plots showing max ΔF/F_0_ values from experiments depicted in (B). (**D**) Example of calcium sensitive fluorescence time-course of cells co-expressing PIEZO1 and GCaMP6f in presence (blue trace) or absence (black trace) of 2.5 μM GsMTx4 and exposed to a shear stress pulse. (**E**) Scatter plots showing max ΔF/F_0_ values obtained from experiments depicted in (D). (**F**) Examples of iGlow fluorescence time course in presence of 2.5 μM GsMTx4. (**G**) Scatter plots showing max ΔF/F_0_ values obtained from (F). (**H**) Representative fluorescence traces from iGlow and H8del. (**I**) Scatter plots showing max ΔF/F_0_ values from experiments shown in (H). (**J**) Histogram comparing time to peak values between iGlow and H8Del from experiments depicted in (H-I). (**K**) Fluorescence time course from H8Del expressing cells obtained with repeated shear stress pulse (1.7 dyne cm^-2^) with 1 min recovery. Insert: amplitude of indicated repeated peaks relative to first peak amplitude (n = 3). Numbers above plots indicate exact p-values from one-way ANOVA (C) or Student’s T-tests (E), (G) and (I-J). Error bars = s.e.m.

Acute incubation with micromolar concentrations of the spider toxin GsMTx4 robustly inhibits a broad range of mechanosensitive ion channels upon various mechanical stimulations, including membrane stretch, fluid shear stress and mechanical indentation^33–37^. Hence, we tested whether this toxin could inhibit iGlow. We first performed control experiments by measuring Ca^2+^ entry mediated by the mechanosensitive Piezo1 channel in response to a 2.6 dyne cm^-2^ pulse in the presence or absence of 2.5 μM GsMTx4. We monitored intracellular free Ca^2+^ ions by co-transfecting PIEZO1-deficient cells^38^ with a mouse Piezo1 plasmid and a plasmid encoding the fluorescent calcium indicator GCaMP6f^39^. Our data show that this toxin concentration was able to reduce GCaMP6fs fluorescence response (max ΔF/F_0_) from +75 ± 5 % to +16 ± 2 %, a nearly 5-fold reduction (Student’s T-test p-value = 9.7 x 10^-18^) (**Figure 6D-E**). In contrast, the same treatment did not affect the amplitude of iGlow signals induced by a 2.6 dyne cm^-2^ pulse (Student’s T-test p-value = 0.9116) (**Figure 6F-G**).

Class-A GPCRs harbor a structurally-conserved amphipathic helical motif located immediately after the seventh transmembrane segment, called Helix 8. A recent study showed that deletion of Helix 8 abolished mechanical, but not ligand-mediated, activation in the histamine receptor H1R^20^. Furthermore, transplantation of H1R Helix 8 into a mechano-insensitive GPCR was sufficient to confer mechanosensitivity to the chimeric receptor^20^. The online tool NetWheels indicates that GPR68 also contains an amphipathic helical motif resembling the Helix 8 of H1R (Supplementary Figure 5). We introduced a non-sense codon (TGA) to eliminate this motif and the remainder of the C-terminal region from iGlow (see Supplementary Figure S1) and tested the sensitivity of the deletion mutant (H8del) to a single FSS pulse of 2.6 dyne cm^-2^. At this shear stress amplitude, the mean peak amplitude produced by H8del was not statistically different than those produced by the full-length iGlow (Student’s T-test p-value = 0.3933) (Figure 4H-I). In addition, the time to peak was similar in both cases (Student’s T-test p-value = 0.8808) (Figure 6J). This result show that Helix 8 is not required for shear flow activation by our sensor. Since the C-terminal deletion eliminates regulatory sites involved in GPCR desensitization, H8Del signals may enable repeated stimulations with less signal loss compared to iGlow. When H8Del was stimulated with a 1.7 dynes cm^-2^ pulse every minute, the peak amplitude remained approximately constant for the first three peaks before fading (Figure 6K).

## Discussion

This study introduces iGlow, a genetically-encoded fluorescent sensor of GPR68 activation. iGlow responds to all currently known activators of GPR68, i.e. fluid shear stress, the synthetic agonist ogerin and extracellular acidifications. A striking observation was the presence of flow-induced activation of iGlow in both intracellular vesicular compartments as well as in membrane-bound structures reminiscent to primary cilia. Since the presence of GPCRs in primary cilia has been reported, it is possible that endogenous GPR68 may reside in endothelial primary cilia to sense shear stress^40,41^. Alkalinization-induced internalization of GPR68 has been recently reported in leukocytes, thus the presence of an intracellular pool of iGlow in fibroblastic HEK293T cells is not surprising^42^. However, internalized GPCRs are commonly considered desensitized and thus unable to activate and/or mediate G protein signaling. In contrast, flow-induced iGlow activation also occurred from intracellular vesicles, suggesting the possibility that GPR68 senses flow from within the cell, and could transduce force into G protein signals. This idea is supported by a growing number of empirical data showing GPCR-mediated signaling from intracellular compartments^43,44^. Further investigations will be necessary to test if internalized GPR68 receptors can mediate flow-induced G-protein signaling. Meanwhile, these results allow us to think about how a dynamic redistribution of GPR68 between the cell surface and subcellular compartments could constitute a cellular mechanism to uncouple proton-dependent from shear stress-dependent activation of GPR68.

Another unexpected property of iGlow is the relatively large cell-to-cell fluctuation of the overall shape and/or peak of its fluorescence signals. These fluctuations may reflect the intrinsic heterogeneity of morphologies and mechanical properties of cultured HEK293T cells which may lead to heterogenous flow-induced cellular deformations and iGlow activation profiles. Another potential contributor of signal fluctuation is the fact that iGlow resides in different cellular compartments. Indeed, the distribution of iGlow in these compartments exhibit large cell-to-cell variations and each compartment may be differentially affected by extracellular shear stress. The heterogeneity of iGlow cellular localizations may also contribute to the fluctuations of iGlow signals induced by extracellular protons and the synthetic agonist ogerin in absence of flow. Despite these fluctuations, the peak amplitude of iGlow fluorescence signals was, on average, larger and occurred quicker when the amplitude of its endogenous stimuli, i.e. shear stress and external protons, was larger.

A main advantage of iGlow is its insensitivity to pharmacological modulation of G-protein signaling, disruption of actin cytoskeleton, and incubation with GsMTx4, a toxin proposed to inhibit mechanosensitive membrane proteins by partitioning into the lipid bilayer, effectively “buffering” the effect of shear stress and/or membrane stretch^45^. As in dLight1.2, the insertion of the cpGFP module near the G-protein binding site likely uncouples iGlow from G-protein signaling. However, the lack of signal elimination by Cytochalasin D and GsMTx4 treatments is more puzzling since numerous mechanosensitive ion channels show at least partial reduction of mechanosensitivity in response to these treatments^33,37,46–52^. The apparent lack of sensitivity of iGlow to Cytochalasin D and GsMTx4 could be, in part, explained by the presence of intracellularly-located iGlow. Indeed, intracellular iGlow would not be inhibited by GsMTx4, a membrane-impermeant proteinogenic toxin, nor by disruption of cortical actin filaments, which are mostly supporting the plasma membrane.

To date, mechanosensitivity has been reported in at least one class-B GPCR (parathyroid hormone type 1 receptor)^53^ and many class-A subfamilies including A3 (bradykinin receptor B2, Apelin receptor and angiotensin II type 1 receptor)^11,54,55^, A6 (vasopressin receptor 1A)^55^, A13 (sphingosine receptor 1)^56^, A15 (GPR68)^9,10^, A17 (dopamine receptor DRD5)^27^ and A18 (muscarinic receptor M5R and histamine receptor H1R)^20,55^. Helix 8 has been shown both necessary and sufficient to confer mechanosensitivity in certain class-A GPCRs such as H1R^20^. However, Helix 8 is not necessary for flow-induced iGlow fluorescence activation. Further investigations will be necessary to identify the molecular mechanism by which GPR68 (and iGlow) senses flow.

To conclude, iGlow robustly probes GPR68 activation by endogenous and exogenous stimuli and will be invaluable to determine the biological roles of GPR68 in vascular and non-vascular physiology, such as hippocampal plasticity, as well as to better understand how mechanical signals are detected and transduced from different cellular and subcellular compartments.

## Methods

### Molecular cloning

A fragment containing the human GPR68 cDNA was obtained by digesting a pBFRT-GPR68 plasmid (a gift from Drs. Mikhail Shapiro, Caltech and Ardèm Patapoutian, Scripps Research) by NdeI and BamHI. The insert was ligated into an in-house pCDNA3.1-Lck-GCaMP6f plasmid linearized by the same enzymes, creating the plasmid pCDNA3.1-GPR68. A cpGFP cassette was amplified by PCR from a pCDNA3.1 plasmid encoding ASAP1 (a gift from Dr. Michael Lin, Stanford, available as Addgene #52519) and inserted into pCDNA3.1-GPR68 using the NEBuilder HiFi DNA Assembly kit (New England Biolabs). The pCNDA3.1-jRGECO1a plasmid was cloned by assembling PCR-amplified fragments from pGP-CMV-NES-jRGECO1a (Addgene # 61563, a gift from Dr. Douglas Kim^57^) and pCDNA3.1. All constructs were confirmed by Sanger sequencing (GENEWIZ). The pLifeAct-mScarlet-N1 plasmid was obtained from Addgene (#85054, a gift from Dr. Dorus Gadella^58^). The DsRed-CAAX and HcRed-CAAX plasmids were a gift from Dr. Bradley Andersen. All molecular biology reagents were purchased from New England Biolabs.

### Cell culture, transfection and drug treatment

HEK293T cells were obtained from the American Tissue Culture Collection and ΔPZ1 cells were a gift from Ardèm Patapoutian (Scripps Research). Cells were cultured in standard conditions (37 °C, 5 % CO_2_) in a Dulbecco’s Modified Eagle’s Medium supplemented with Penicillin (100 U mL^-1^), streptomycin (0.1 mg mL^-1^), 10 % sterile Fetal Bovine Serum, 1X Minimum Essential Medium non-essential amino-acids and without L-glutamine. All cell culture products were purchased from Sigma-Aldrich. Plasmids were transfected in cells (passage number < 35) seeded in 96-well plates at ~50 % confluence 2-4 days before the experiment with FuGene6 (Promega) or Lipofectamine 2000 (Thermo Fisher Scientific) and following the manufacturer’s instructions. 1-2 days before experiments, cells were gently detached by 5 min incubation with Phosphate Buffer Saline and re-seeded onto 18 mm round glass coverslips (Warner Instruments) or onto disposable flow chambers (Ibidi μ-slides 0.8 or 0.4mm height), both coated with Matrigel (Corning). Cells were treated with each drug at 15 min (CMPD101), 20 min (cytochalasin D and NF449), 30 min (GTP-gamma-S and GsMTx4) or 2 hrs (BIM-46187) prior to measurement. pH-shear experiments were performed using a starting pH of 8.2, adjusted 15 min prior to measurement. CMPD101 (#5642) and NF-449 (#1391) were purchased from R&D Systems (Biotechne), GTP-gamma-S was purchased from Cytoskeleton, Inc (#BS01), Dopamine (#H8502) and Gαq inhibitor BIM-46187 (#5332990001) were purchased from Sigma-Aldrich.

### Fluorescence imaging

Excitation light of desired wavelengths were produced by a Light Emitting Diode light engine (Spectra X, Lumencor), cleaned through individual single-band excitation filters (Semrock) and sent to the illumination port of an inverted fluorescence microscope (IX73, Olympus) by a liquid guide light. Excitation light was reflected towards a plan super apochromatic 100X oil-immersion objective with a 1.4 numerical aperture (Olympus) using a triple-band dichroic mirror (FF403/497/574, Semrock). Emission light from the sample was filtered through a triple-band emission filter (FF01-433/517/613, Semrock) and sent through beam-splitting optics (W-View Gemini, Hamamatsu). Split and unsplit fluorescence images were collected by a sCMOS camera (Zyla 4.2, ANDOR, Oxford Instruments). Spectral separation by the Gemini was done using flat imaging dichroic mirrors and appropriate emission filters (Semrock). Images were collected by the Solis software (ANDOR, Oxford Instruments) at a rate of 1 frame s^-1^ through a 10-tap camera link computer interface. Image acquisition and sample illumination were synchronized using TTL triggers digitally generated by the Clampex software (Molecular Devices). To reduce light-induced bleaching, samples were only illuminated for 200 ms, i.e. during frame acquisition (200 ms exposure). To reduce auto-fluorescence, the cell culture medium was replaced with phenol red-free HBSS approximately 20 min prior experiments. Due to the narrow field of view at this magnification, only 1-3 transfected cells were measured per each flow assay.

### Image analyses

A MATLAB script (**Dataset S1**) was used to calculate the average fluorescence intensity of each cell of interest at each frame, expressed as percentile above the fluorescence at t = 0 (“deltaF/F”). Prior to analysis, photobleaching was corrected using the exponential fit method and a “mask” (.jpg format) separating cells of interest from the background was manually drawn in ImageJ. This mask, in addition to a bleach-corrected image sequence (.tif format) corresponding to each frame of the video, is read by the MATLAB script. The output of the script is a text file containing “deltaF/F” values at each frame for each cell in the mask (“bleachcorrected.txt”, rows and columns correspond to frame ID and individual cells respectively), and a figure showing the original image, the mask, and deltaF/F traces for each cell (“figure.jpg”). Backgrounds are either removed manually in ImageJ, or within the MATLAB script by taking the average fluorescence intensity of all non-cell pixels and subtracting this value from the intensity values associated with cells at each frame.

### Fluid shear stress stimulation and calculations

Fluid shear-stress stimulation was done by circulating extracellular physiological solutions at various speeds into a μ-slide channel (Ibidi) using a Clampex-controlled peristaltic pump (Golander). The average amplitude of wall shear stress *τ* applied at the cell surface was estimated using the manufacturer’s empirical equation relating τ with flow rate Φ for μ-slide channel with 0.4 mm height:

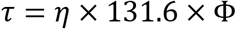

We independently measured *τ* using particle image velocimetry measurements. Briefly, we dispensed 6 μm diameter polystyrene beads (Polybead microspheres, Polysciences) diluted in HBSS into recording μ-slide channels and let them settle to the bottom of the μ-slide. Beads were imaged using a 100X immersion objective (Olympus) focused at the fluid-wall boundary. Bead velocity was estimated using high-speed imaging (300-500 frames s^-1^) for various flow rates (Supplementary Table 1). τ depends on the distance between the fluid and the boundary *y*, the dynamic viscosity of the fluid *μ* and the flow velocity *u* according to:

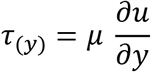

Since the beads are located very close to the boundary, we can assume they are within the viscous sublayer^59^. Hence, in these conditions, the fluid velocity profile is linear with the distance from the boundary:

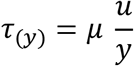

Shear stress values were calculated using the experimentally measured *u* values (in m s^-1^) and using an averaged bead radius of 2.5 x 10^-6^ m. For *μ*, we measured the dynamic viscosity of HBSS at room temperature using a rotary viscometer (USS-DVT4, U.S. SOLID) and obtained a value of 1.04 ± 0.04 x 10^-3^ Pa s^-1^ (n = 3).

### Statistical analyses

The number n represents the number of cells analyzed. To evaluate pairwise differences of mean data sets, we performed Mann-Whitney U-tests when n ≤ 10 and Student’s T-tests when n > 10 in both data sets. To compare difference of mean values between different treatments, we performed one-way ANOVA. All error bars are standard errors of the mean. All statistical tests were performed on OriginPro 2018.

## Data availability

Data obtained from fluorescence traces (ΔF/F_0_ and time to peak values) have been deposited in the Open Science Framework (OSF) public Depository (DOI 10.17605/OSF.IO/8MP4W).

## Acknowledgments

We thank Ardèm Patapoutian for the gift of the human GPR68 cDNA and Bradley Andresen for help with confocal imaging. This work was supported by intramural and start-up funds from Western University of Health Sciences (to J.J.L), federal work-study (to T.G) and NIH grants GM130834 and NS101384 (to J.J.L.).

## Author contributions

J.J.L. conceived the project. A.D.O., T.G., A.T. and W.P. performed experiments. A.D.O., T.G. and J.J.L. analyzed data. J.J.L. wrote the manuscript with input from A. D. O.

