## supplementary information for "Mechanical and chemical activation of GPR68 (OGR1) probed with a genetically-encoded fluorescent reporter"

**This PDF file includes:**

Figures S1 to S5

Table S1

ATGGGGAACATCACTGCAGACAACTCCTCGATGAGCTGTACCATCGACCATACCATCCACCAGACGCTGGCCCCGGTGGTCTATGTTACCGTGCTGGTGGTGGGCTTCCCGGCCAACTGCCTGTCCCTCTACTTCGGCTACCTGCAGATCAAGGCCCGGAACGAGCTGGGCGTGTACCTGTGCAACCTGACGGTGGCCGACCTCTTCTACATCTGCTCGCTGCCCTTCTGGCTGCAGTACGTGCTGCAGCACGACAACTGGTCTCACGGCGACCTGTCCTGCCAGGTGTGCGGCATCCTCCTGTACGAGAACATCTACATCAGCGTGGGCTTCCTCTGCTGCATCTCCGTGGACCGCTACCTGGCTGTGGCCCATCCCTTCCGCTTCCACCAGTTCCGGACCCTGAAGGCGGCCGTCGGCGTCAGCGTGGTCATCTGGGCCAAGGAGCTGCTGACCAGCATCTACTTCCTGATGCACGAGGAGGTCATCGAGGACGAGAACCAGCACCGCGTGTGCTTTGAGCACTACCCCATCCAGGCATGGCAGCGCGCCATCAACTACTACCGCTTCCTGGTGGGCTTCCTCTTCCCCATCTGCCTGCTGCTGGCGTCCTACCAGGGCATCCTGCGCGCCGTGCGCCGGAGCCTGAGCTCACTCATTAACGTCTATATCAAGGCCGACAAGCAGAAGAACGGCATCAAGGCGAACTTCAAGATCCGCCACAACATCGAGGACGGCGGCGTGCAGCTCGCCTACCACTACCAGCAGAACACCCCCATCGGCGACGGCCCCGTGCTGCTGCCCGACAACCACTACCTGAGCGTGCAGTCCAAACTTTCGAAAGACCCCAACGAGAAGCGCGATCACATGGTCCTGCTGGAGTTCGTGACCGCCGCCGGGATCACTCTCGGCATGGACGAGCTGTACAAGGGCGGTACCGGAGGGAGCATGGTGAGCAAGGGCGAGGAGCTGTTCACCGGGGTGGTGCCCATCCTGGTCGAGCTGGACGGCGACGTAAACGGCCACAAGTTCAGCGTGTCCGGCGAGGGTGAGGGCGATGCCACCTACGGCAAGCTGACCCTGAAGTTCATCTGCACCACCGGCAAGCTGCCCGTGCCCTGGCCCACCCTCGTGACCACCCTGACCTACGGCGTGCAGTGCTTCAGCCGCTACCCCGACCACATGAAGCAGCACGACTTCTTCAAGTCCGCCATGCCCGAAGGCTACATCCAGGAGCGCACCATCTTCTTCAAGGACGACGGCAACTACAAGACCCGCGCCGAGGTGAAGTTCGAGGGCGACACCCTGGTGAACCGCATCGAGCTGAAGGGCATCGACTTCAAGGAGGACGGCAACATCCTGGGGCACAAGCTGGAGTACAACAATCATGACCAACTGAGCCGCAAGGACCAGATCCAGCGGCTGGTGCTCAGCACCGTGGTCATCTTCCTGGCCTGCTTCCTGCCCTACCACGTGTTGCTGCTGGTGCGCAGCGTCTGGGAGGCCAGCTGCGACTTCGCCAAGGGCGTTTTCAACGCCTACCACTTCTCCCTCCTGCTCACCAGCTTCAACTGCGTCGCCGACCCCGTGCTCTACTGCTTCGTCAGCGAGACCACCCACCGGGACCTGGCCCGCCTCCGCGGGGCCTGCCTGGCCTTCCTCACCTGCTCCAGGACCGGCCGGGCCAGGGAGGCCTACCCGCTGGGTGCCCCCGAGGCCTCCGGGAAAAGCGGGGCCCAGGGTGAGGAGCCCGAGCTGTTGACCAAGCTCCACCCGGCCTTCCAGACCCCTAACTCGCCAGGGTCGGGCGGGTTCCCCACGGGCAGG

MGNITADNSSMSCTIDHTIHQTLAPVVYVTVLVVGFPANCLSLYFGYLQIKARNELGVYLCNLTVADLFYICSLPFWLQYVLQHDNWSHGDLSCQVCGILLYENIYISVGFLCCISVDRYLAVAHPFRFHQFRTLKAAVGVSVVIWAKELLTSIYFLMHEEVIEDENQHRVCFEHYPIQAWQRAINYYRFLVGFLFPICLLLASYQGILRAVRRSLSSLINVYIKADKQKNGIKANFKIRHNIEDGGVQLAYHYQQNTPIGDGPVLLPDNHYLSVQSKLSKDPNEKRDHMVLLEFVTAAGITLGMDELYKGGTGGSMVSKGEELFTGVVPILVELDGDVNGHKFSVSGEGEGDATYGKLTLKFICTTGKLPVPWPTLVTTLTYGVQCFSRYPDHMKQHDFFKSAMPEGYIQERTIFFKDDGNYKTRAEVKFEGDTLVNRIELKGIDFKEDGNILGHKLEYNNHDQLSRKDQIQRLVLSTVVIFLACFLPYHVLLLVRSVWEASCDFAKGVFNAYHFSLLLTSFNCVADPVLYCF**[**VSETTHRDLARLRGACLAFLTCSRTGRAREAYPLGAPEASGKSGAQGEEPELLTKLHPAFQTPNSPGSGGFPTGR**]**

**Fig. S1. Nucleic acid (top) and amino acids (bottom) sequences of iGlow.** Black = GPR68; purple = linkers; green = cpGFP; red = Helix 8. Brackets indicate the C-terminal fragment eliminated in H8Del.

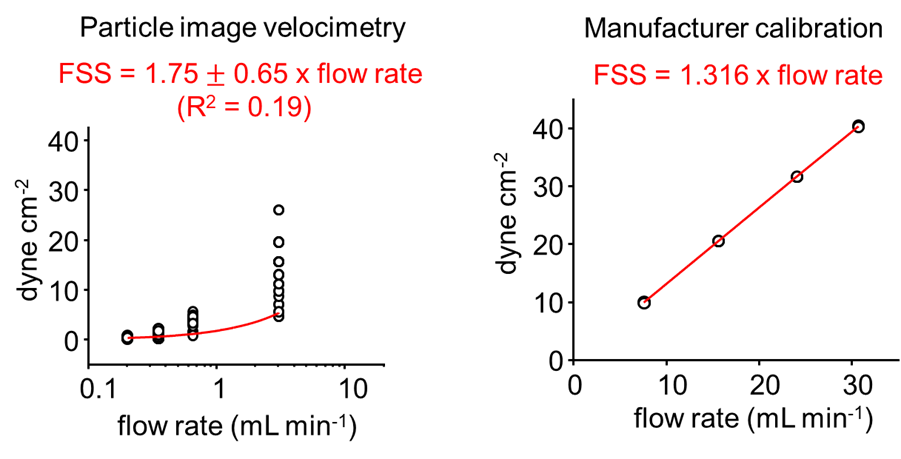

**Fig. S2.** **Shear stress calibration.** Shear stress applied through our flow chamber was calculated using particle image velocimetry (*left*, see methods) or using the manufacturer's calibration (*right*).

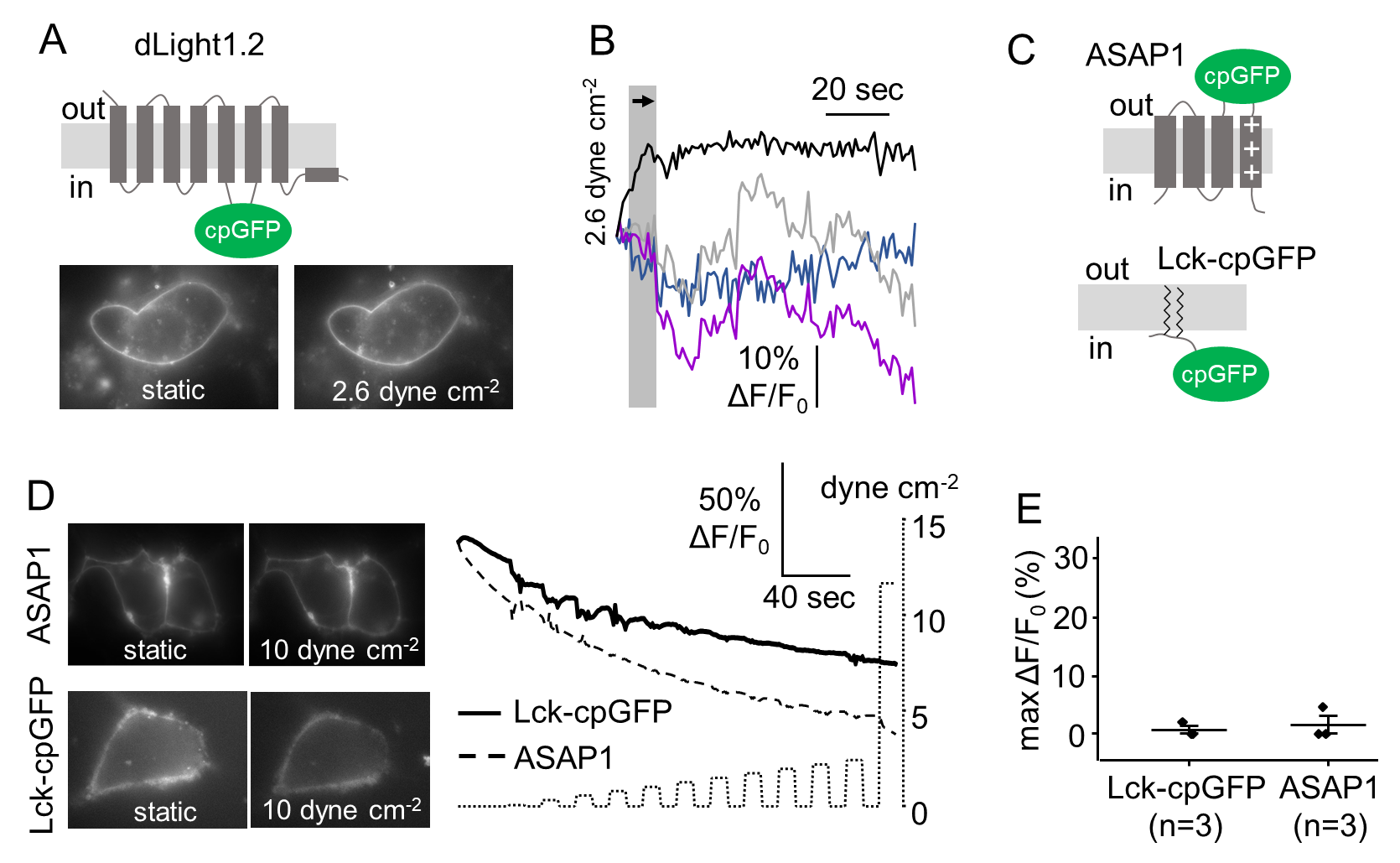

**Fig. S3.** **cpGFP is not directly modulated by shear stress.** (**A**) *Top*: cartoon representing the position of cpGFP in dLight1.2. *Bottom*: epifluorescence images of cells expressing dLight1.2 under static or flow conditions. (**B**) Representative fluorescence time courses of dLight1.2 in response to the indicated shear stress pulse. (**C**) Cartoons showing the position of cpGFP in ASAP1 and Lck-cpGFP. (**D**) *Left*: epifluorescence images of cells expressing ASAP1 (top) or Lck-cpGFP (bottom) under static of flow conditions. *Right*: example of fluorescence time-courses from cells expressing ASAP1 (dashed line) or Lck-cpGFP (solid line) in response to FSS pulses of incrementally increased amplitudes (dotted line). (**E**) Scatter plots showing the maximal ΔF/F_0_ values obtained with ASAP1 (n = 3) and Lck-cpGFP (n = 3) using the FSS protocol shown in (B).

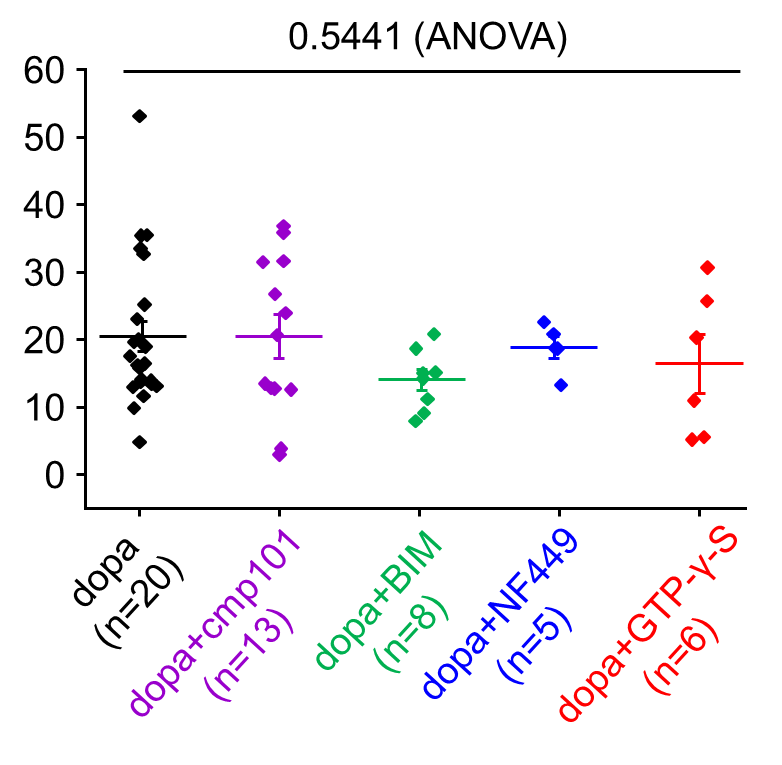

**Fig. S4.** **Dopamine sensitivity of dLight1.2 is not abrogated by pharmacological modulation of G protein signaling.** The scatter plots show max ΔF/F_0_ values obtained upon acute perfusion with 10 µM dopamine in cells pre-treated with CMPD101 (purple dots, n = 13), BIM-46187 (BIM, green dots, n = 8), NF-449 (blue dots, n = 5), GTP-γ-S (red dots, n = 6), or a vehicle control (black dots, n = 20). The number above the graph indicates the exact p-value from a one-way ANOVA. Error bars = s.e.m.

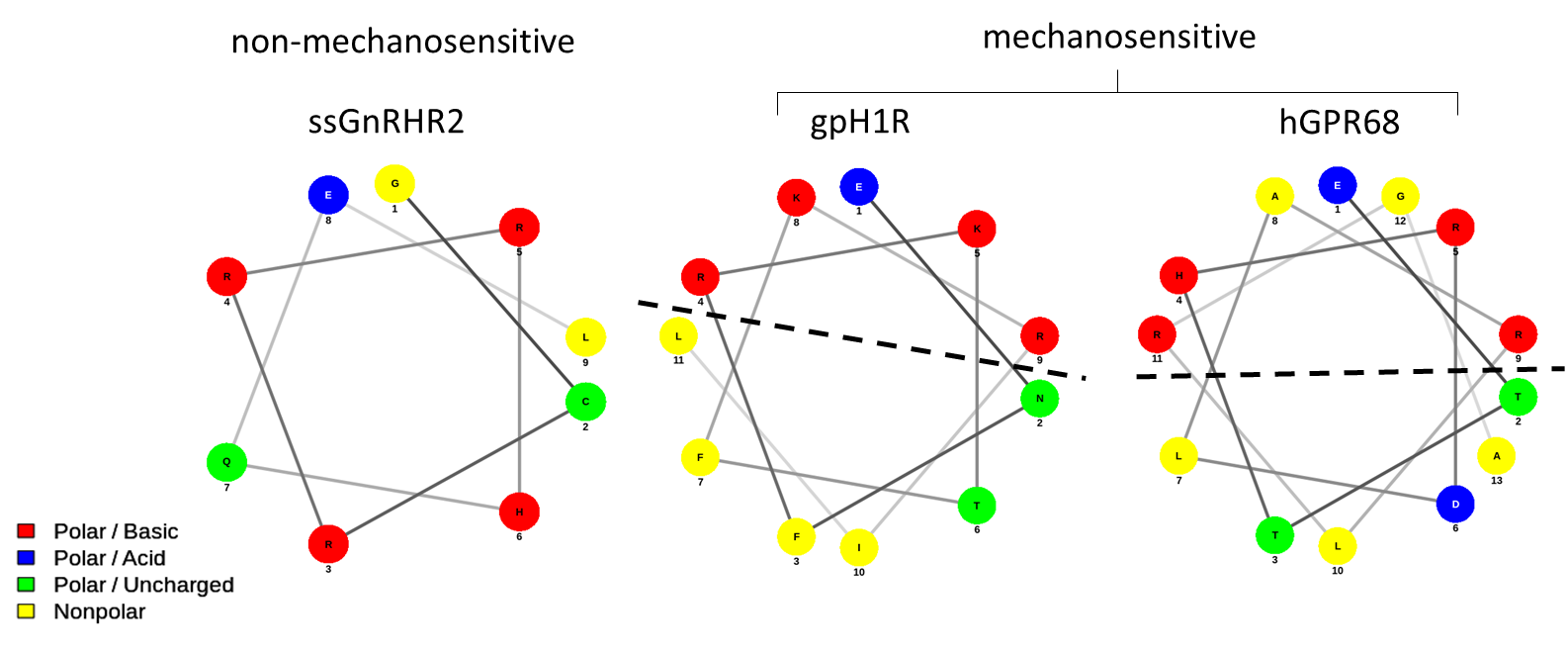

**Fig. S5.** **Prediction of an amphipathic Helix 8 in GPR68 using the online predictor NetWheels.** Helical wheel plot of Helix 8 in the long isoform of the swine gonadotropin-releasing hormone receptor (ssGnRHR2), the guinea pig histamine H1 receptor (gpH1R) and human GPR68 (hGPR68). The dotted line indicates the separation between the polar vs. apolar interfaces of the Helix.

| **primers** | **Sequences (5'→3')** |
| --- | --- |
| GPR68 Fwd (backbone) | aatcatgaccaactgagccgcaaggaccagatccagcgg |
| GPR68 Rev (backbone) | aatgagtgagctcaggctccggcgcacggcgcg |
| cpGFP with linkers Fwd | gcgccggagcctgagctcactcattaacgtctatatcaaggcc |
| cpGFP with linkers Rev | ccttgcggctcagttggtcatgattgttgtactccagcttgtg |

**Table S1. Primers used to insert cpGFP into GPR68 using DNA assembly.**
