## Supplementary material for "Mechanical and chemical activation of GPR68 (OGR1) probed with a genetically-encoded fluorescent reporter": dataset1

**Dataset S1**

**MATLAB script**

```
% Identifies the directory containing the files of interest. This script
% requires a series of sequentially named image files (.tif format,
% typically obtained by exporting a video file as image series in ImageJ)
% and an accompanying 'mask' image containing only the regions of interest,
% with other sections cropped out (.jpg format, obtained through the 'clear
% outside' command in ImageJ). Each set of images must be contained in its
% own folder.
cd 'K:\test'
D = dir;
% Gets a list of folders within the directory and enters each to perform
% deltaF/F analysis in a loop.
for k = 3:length(D);
    currD = D(k).name;
    cd (currD);
% Finds the image series in each folder and determines the number of frames.
% In this case, images are labeled 'sample-0000.tif' through
% 'sample-0096.tif' in a 97-frame video.
    c = struct2cell(dir('*.tif'));
        firstname = c{2,1}
        secondname = c{1,2}
        combinedname = [firstname,'\',secondname];
        countimages = dir('*.tif');
        lastimageid = length(countimages);
        lastimageid = lastimageid - 1;
% Identifies and reads the first file in the series ('sample-0000.tif').
        firstimageid = 0
        imageidstr = num2str(firstimageid);
        basename = combinedname(1:end-5);
        initialname = [basename,imageidstr,'.tif'];
        I = imread(initialname)
 % Enhances contrast and removes background noise using a Wiener filter.
        I = adapthisteq(I);
        I = wiener2(I, [5 5]);
        framelimit = lastimageid;
 % Identifies the 'mask' image in the folder.
        m = struct2cell(dir('*.jpg'));
            firstname_m = m{2,1}
            secondname_m = m{1,1}
            combinedname_m = [firstname_m,'\',secondname_m];
            MaskImage = imread(combinedname_m);
 % Uses Otsu's method to set a threshold for segmenting the image. This
 % process is error-prone in complex images, and masks are used in the
 % present analysis to ensure that the segmentation is faithful to cell
 % boundaries.
        threshold = graythresh(MaskImage);
        bw = im2bw(MaskImage, threshold);
 % Combines gaps in the mask and removes any segment that is less than
 % 10000 pixels in area. This area threshold is suitable for
 % high-magnification images (100x), images under lower magnification must
 % use lower thresholds to adjust for the smaller areas associated with
 % cells.
        bw2 = imfill(bw,'holes');
        bw3 = bwareaopen(bw2, 10000);
        bw3_perim = bwperim(bw3);
 % Creates a second perimeter with the inverse of the mask, to be used for
 % background correction.
        bwinvert = imcomplement(bw3);
 % Creates an overlay of the first image of the series and the regions of
 % interest as determined by the mask and segmentation.
        overlay1 = imoverlay(I, bw3_perim, [1 .3 .3]);
        L = bwlabel(bw3);
        s = regionprops(L, 'Centroid');
 % Adds the overlay image to the figure, to be saved to each folder later.
        figure
        plot1 = subplot (2,1,1);
        imshow(overlay1)
        [columns,rows] = size(s(:,1));
        minz0=zeros(columns,framelimit);
        intensityfull=minz0;
        backgroundsraw=minz0;
        hold on
        for k = 1:numel(s)
            c = s(k).Centroid;
            text(c(1), c(2), sprintf('%d', k), 'HorizontalAlignment', 'center', 'VerticalAlignment', 'middle', 'color', 'g');
        end
        hold off
 % Identifies the shared name between image files. The deletion of eight
 % characters at the end of the file ID ensures that series with up to 1000
 % frames can be read by the script. In this case, the shared name would be
 % the directory address of the file series, followed by 'sample-'.
        basename = combinedname(1:end-8);
 % For each frame in the series, finds the frame number and converts this
 % to a 4-character string to be appended to the end of the shared name.
 % For example, the 30th frame would yield a string of '0029' and a file
 % name of 'sample-0029.tif'.
        try
        for imid = 1:lastimageid;
            imstr=num2str(imid);
            while length(imstr) < 4;
                imstr = strcat('0',imstr);
            end
            filename = [basename,imstr,'.tif'];
 % Reads the frame, determines the raw fluorescence intensity at each cell
 % of interest in each frame, and adds this to an array. At the same time,
 % reads the raw fluorescence intensity in non-cell regions and adds this
 % to a second array.
            Ianalysis = imread(filename);
            intensities = regionprops(bw3, Ianalysis, 'MeanIntensity');
            backgroundintensities = regionprops(bwinvert, Ianalysis, 'MeanIntensity');
            fullid = imid;
            A = struct2cell(intensities);
            B = struct2cell(backgroundintensities);
            out = cat(2,A{:});
            intensityfull(1:columns,fullid)=out
            out2 = cat(2,B{:});
            backgroundsraw(1:columns,fullid)=out2
        end;
        final = intensityfull';
        backgroundsfinal = backgroundsraw';
 % Performs background correction by subtracting the background intensity
 % from the cellular fluorescence intensity at each frame. This step should
 % be skipped if the backgrounds were already corrected using ImageJ.
        final = bsxfun(@minus,final,backgroundsfinal);
 % Determines the deltaF/F0 values.
        deltaF = bsxfun(@minus,final,final(1,:));
        bleachcorrected = bsxfun(@rdivide,deltaF,final(1,:));
 % Adds deltaF/F traces to the figure.
        plot2 = subplot (2,1,2);
        plot(bleachcorrected)
        set(gca,'XLim',[0 lastimageid])
        ylabel('Fold change in intensity');
        xlabel('Frame number');
        title('\rmIntensity fluctuations');
 % Saves the deltaF/F values as 'bleachcorrected.txt' and the figure as
 % 'figure.jpg'.
        save ('bleachcorrected.txt','bleachcorrected','-ascii')
        saveas (gcf,'figure.jpg');
close all;
        end
cd ('..');
   end;
```
